# Mix-and-extrude: high-viscosity sample injection towards time-resolved protein crystallography

**DOI:** 10.1101/2022.11.23.517685

**Authors:** Mohammad Vakili, Huijong Han, Christina Schmidt, Agnieszka Wrona, Marco Kloos, Iñaki de Diego, Katerina Dörner, Tian Geng, Chan Kim, Faisal Koua, Diogo Melo, Mathieu Rappas, Adam Round, Ekaterina Round, Marcin Sikorski, Joana Valerio, Tiankun Zhou, Kristina Lorenzen, Joachim Schulz

## Abstract

Time-resolved crystallography enabled the visualization of protein molecular motion during reaction. While light is commonly used to initiate reactions in time-resolved crystallography, only a small number of proteins can in fact be activated by light. However, many biological reactions can be triggered by the interaction of proteins with ligands. The sample delivery method presented here uses a mix-and-extrude approach based on 3D printed microchannels in conjunction with a micronozzle to study the dynamics of samples in viscous media that can be triggered by diffusive mixing. The device design allows for mixing of ligands and protein crystals in a time window of 2 to 20 seconds. The device characterization using a model system (fluorescence quenching of iq-mEmerald proteins by copper ions) demonstrated that ligand and protein crystals, each within the lipidic cubic phase, can be mixed efficiently. The potential use of this approach for time-resolved membrane protein crystallography to support in the development of new drugs is also discussed.

**Synopsis:** 3D printed mixing-HVE devices address time-resolved membrane protein crystallography challenges via compact dual-flow LCP injection.

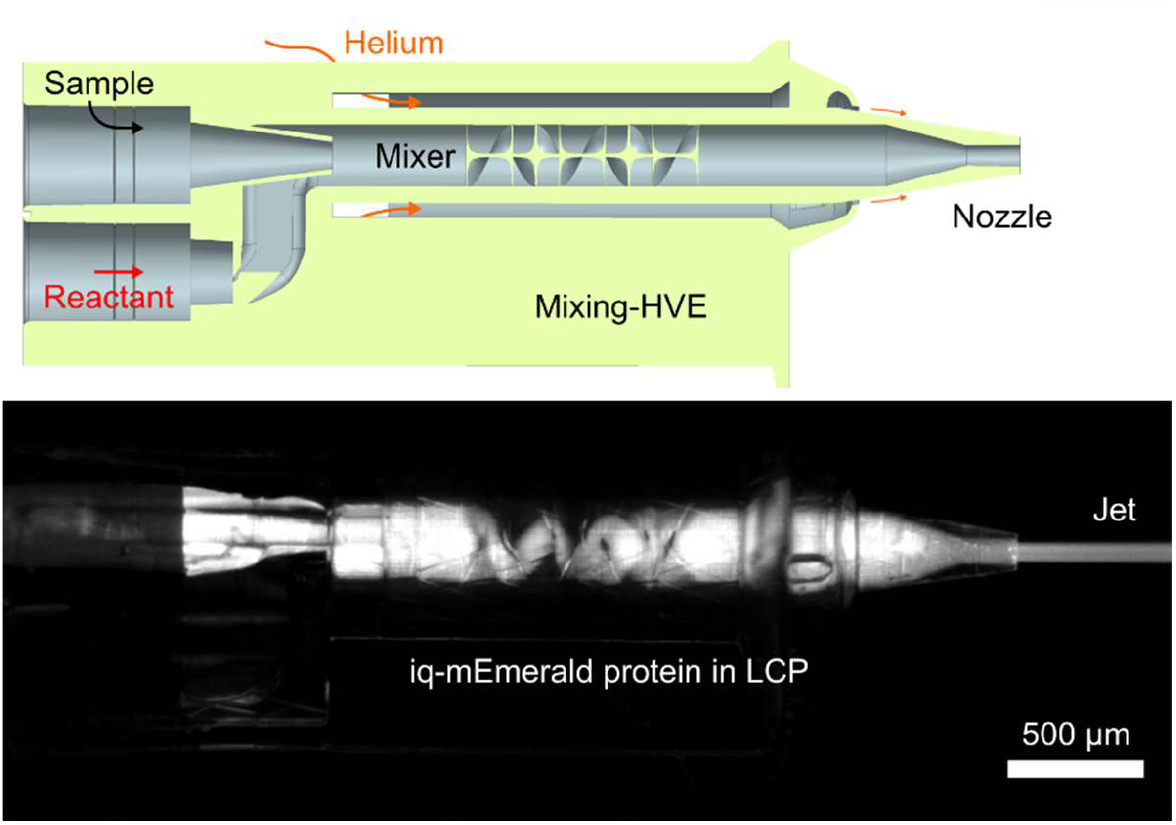

## 1. Introduction

Membrane proteins comprise about 23% of all proteins (Uhlén *et al*., 2015). They play an essential role in biological function and are the target of numerous medications on the market (Overington *et al*., 2006).

Due to sample preparation challenges, the number of membrane protein structures that have been determined is rather low despite their biological significance. For rational drug design, the structures of target protein of apo and ligand-bound forms are essential to understand the mode of action.

By employing lipidic cubic phase (LCP) as a medium, membrane protein crystallization was facilitated (Landau & Rosenbusch, 1996; Li & Caffrey, 2020). The LCP features a unique lipid bilayer and aqueous channel structure that is continuous, folded, and highly curved. During crystallization, the cubic phase is locally converted to the lamellar phase by equilibrating with precipitant solutions, and the protein is concentrated in the lamellar phase to form a nucleus and ultimately a crystal (Cherezov & Caffrey, 2007; Caffrey, 2008). Multiple characterization studies demonstrated the mobility of chemicals and proteins in LCP via lipidic or aqueous phases (Cherezov *et al*., 2006; Li *et al*., 2017; Li & Caffrey, 2011; Clogston & Caffrey, 2005; Eriksson & Lindblom, 1993; Boland *et al*., 2018), indicating that LCP can be used as a medium for biochemical/biophysical characterization in addition to crystallization of membrane proteins.

Sample delivery for time-resolved serial crystallography with light-sensitive protein crystals is not different from the ones for serial crystallography for static structure determination; the only difference in the set-up is the additional light source, often a laser, to initiate the reaction. Time-resolved studies of ligand binding, on the other hand, need an additional liquid channel for the ligand. For crystals grown in aqueous solution, mix-and-inject schemes using liquid jets (Pandey *et al*., 2021; Hejazian *et al*., 2020; Calvey *et al*., 2016) or adding/injecting ligand on top of crystals for fixed target, drop-on-demand and tape-drive have been developed as sample delivery for time-resolved serial crystallography (Mehrabi *et al*., 2019; Butryn *et al*., 2021; Beyerlein *et al*., 2017).

The high-viscosity extruder (HVE), created by Weierstall *et al*., is one of the most widely used sample delivery methods for membrane protein crystals in serial crystallography (Weierstall *et al*., 2014). Within this HVE, a hydraulic plunger is used to amplify the pressure provided by an HPLC pump 14 times. This can provide the pressure up to 10,000 psi to drive the sample from the sample reservoir into the capillary and the nozzle. Compared with glass syringes, which can withstand pressures of up to 1,000 psi, this type of injection is more reliable for delivering viscous samples (Grünbein & Kovacs, 2019).

Three-dimensional (3D) printing using two-photon polymerization (2PP) enables a rapid, reproducible and high-throughput nozzle fabrication, in addition to design flexibility (Knoška *et al*., 2020; Nelson *et al*., 2016). Recently, we presented our portfolio of 3D printed sample delivery devices, including viscous extrusion tips that provide controllable sample streams (Vakili *et al*., 2022) and are fully compatible with Weierstall and co-workers’ well established HVE injection hardware. Here, we introduce the 3D printed mix-and-extrude device, which was designed for the simultaneous use of two HVE setups. The device provides a second capillary port for introducing a ligand dispersed in viscous medium. With this, the mixing of two samples immediately before X-ray probing can be achieved. Thus, the mixing-HVE has a great potential in enabling the time-resolved crystallographic study of membrane proteins for the investigation of ligand-binding processes.

Besides their herein reported usage at an FEL, our devices are also suited to the millisecond exposures used at synchrotrons. As our 3D printed nozzles (both with and without mixing capabilities) are fully compatible with Weierstall’s HVE injector - they use the same standard 1/16 inch OD steel tubings as connective part - and the fact that our flow velocity regime (0.25-5.0 mm s^-1^) for stable viscous extrusion matches the parameters from previous serial synchrotron crystallography experiment (Botha *et al*., 2015; Nogly *et al*., 2015; Weinert *et al*., 2017; Botha *et al*., 2018), they represent promising alternatives to the conventionally utilized ground glass capillary tips, that are prone to manual error and lack device reproducibility.

## 2. Materials and Methods

### 2.1. Design choices

The mixer provides a dual-inlet section accepting two capillaries and allowing the convergence of two fluid channels via overlapping concentric cones (Fig. 1A). At the start of the mixing channel, the main/side channel diameter ratio is 100:231.7 μm. Due to the centered sample inlet and the 3D hydrodynamic flow-focusing geometry, a wall contact of the sample is initially (throughout 500 μm in downstream direction) prevented. The total length of the mixing channel is 2570 μm (design “J_7”). From this length, the initial 2070 μm have a 231.7 μm channel width, followed by a 300 μm long tapering section (truncated cone which reduces the ID down to 75 μm), leading to a 200 μm long final section with a 75 μm inner diameter. For the geometry at the tip, the ID-OD-D (liquid channel diameter-gas orifice-distance between orifices) are chosen to be 75-345-600 [μm], therefore, 75 μm wide streams are provided for the X-ray beam.

**Figure 1.**
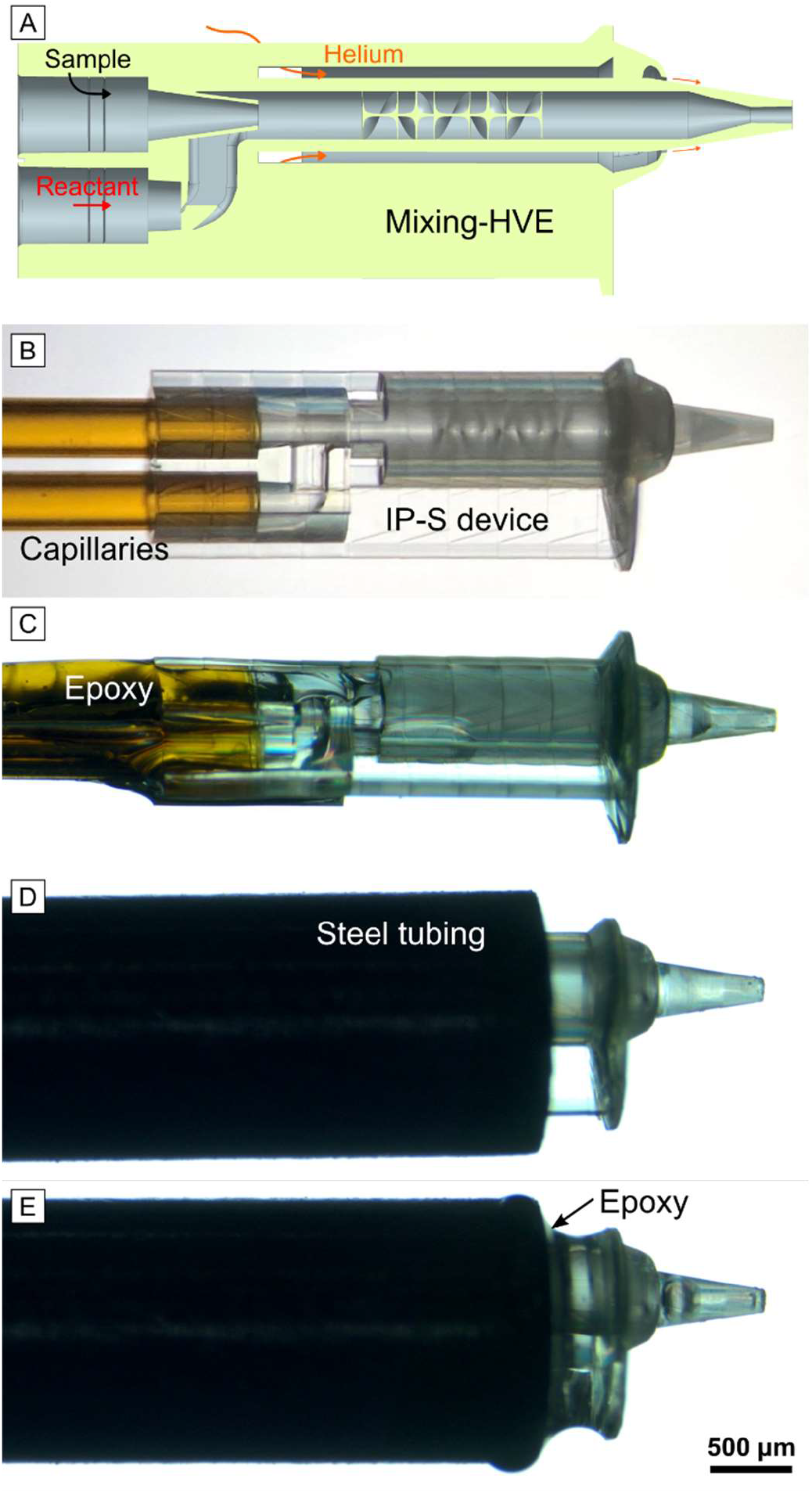
(A) Cross-sectional schematic depiction of the 2PP-3D printed mix-and-extrude device. (B-E) Microscopy images showing the assembly of the mixing-HVE tip with two fluid-feeding fused silica capillaries. (B) Two capillaries, each with 250 μm ID, are inserted into the 3D printed mixing-HVE’s access ports and (C) glued with epoxy glue. (D) The capillaries (un-glued end) are fiddled through a 0.046’’ ID steel tubing (OD = 1/16’’), which is pulled over the mixing section of the device. (E) A small amount of slightly viscous epoxy glue (cured for 2 min) is added onto the gap with a clean capillary or metal wire to secure IP-S to the steel and provide gas tightness.

Inside the first section of the 231.7 μm wide mixing channel, a modified mixing structure based on the “JKMH#10” Kenics mixer from Knoška and Heymann (Knoška *et al*., 2020), is incorporated. With this, a series of six helical elements are introduced into the mixing channel for repeated flow splitting/stretching. The first blade is positioned 500 μm after the mixing initiation point (overlap of the liquid apertures).

With the 2570 μm long mixing channel and a combined liquid flow rate of *Q*_total_= 1.46 μL min^-1^ (*i*.*e*. 1.43 μL min^-1^ for the reactant and 0.03 μL min^-1^ for the sample), a retention time of 4.1 s before extrusion can be achieved within the 3D printed part. This corresponds to a stream velocity of *v*_flow_ = 5.5 mm s^-1^ for the 75 μm wide sample stream. Longer retention times require lower flow rates. For instance, as used during the diffraction data collection described below, a retention time of 18.2 s can be obtained with *Q*_total_ = 0.36 μL min^-1^ (exposed sample has a flow velocity *v*_flow_ = 1.3 mm s^-1^). Another design variation (“J_8”) contains a shorter mixing channel (total length: 1685 μm). With the aforementioned flow rates, this shorter mixing-HVE allows retention times between 2.3 s and 9.9 s.

All 3D CAD files can be found in our online design repository: https://github.com/flmiot/EuXFEL-designs.

At the European XFEL, the X-ray pulses can arrive at 10 Hz - *i*.*e*. one pulse per train. From previous studies, it is known that flow velocities as low as *v* = 0.3 mm s^-1^ are sufficient for stable viscous sample extrusion while maintaining a pulse displacement of *ca*. 30 μm onto the sample and as a result effectively avoiding the exposure of the same crystal with multiple X-ray pulses (Vakili *et al*., 2022). Moreover, sample delivery to X-rays that arrive at higher pulse repetition rates, as seen at the FELs SACLA (60 Hz), PAL-XFEL (60 Hz), SwissFEL (100 Hz) and LCLS (120 Hz), can be accommodated as well (Shimazu *et al*., 2019; Lee *et al*., 2020; Milne *et al*., 2017; Wells *et al*., 2022). However, for ≥100 Hz operation, a sample width reduction from 75 to 50 μm is highly recommended. At constant flow rates, this will increase the flow velocity by a factor or 2.2 that in turn facilitates a viscous extrusion without the need to experience higher pump pressures.

### 2.2. Beamtime injection setup

The SPB/SFX instrument of the European XFEL comprises two interaction regions, a high-vacuum upstream sample environment (Interaction Region Upstream, IRU) and an in-helium downstream interaction region (Interaction Region Downstream, IRD) with respect to the X-ray beam. The instrument setup at IRD provides high flexibility on sample deliveries, including the HVE injection setup. The injection rod funnel attached helium atmosphere sample chamber is located between vacuum out-coupling acoustic delay line (ADL) and detector (JUNGFRAU 4M), and the inserted nozzle rod is positioned using XYZ motors of the goniometer support tower (Round *et al*., in preparation).

The assembled nozzle, connected to capillaries and the 1/16’’ OD steel tubing (IDEX, part no. U-145), was connected to the nozzle holder using a F333N fitting (IDEX) (Weierstall, 2014). A slit on the insertion rod (Fig. S1B) was created 10 cm above the o-ring to shorten the sample capillary length. After screwing the nozzle holder onto the metal rod and inserting the rod into the IRD sample chamber, the capillaries were carefully taken through the slit and linked to the sample reservoirs of the HVEs using 0.015-0.0625” ID PEEK (polyether ether ketone) tubing and custom #10-32 UNF steel fitting with 1/16’’ through-hole (similar to part no. F-354, IDEX). Using ThorLab ½” mounting rods, the two HVE systems were mounted onto the insertion rod to bridge the short distance to the nozzle (Fig. 2A). The final capillary length was 30 cm, with an interior volume of 14.7 μL.

**Figure 2.**
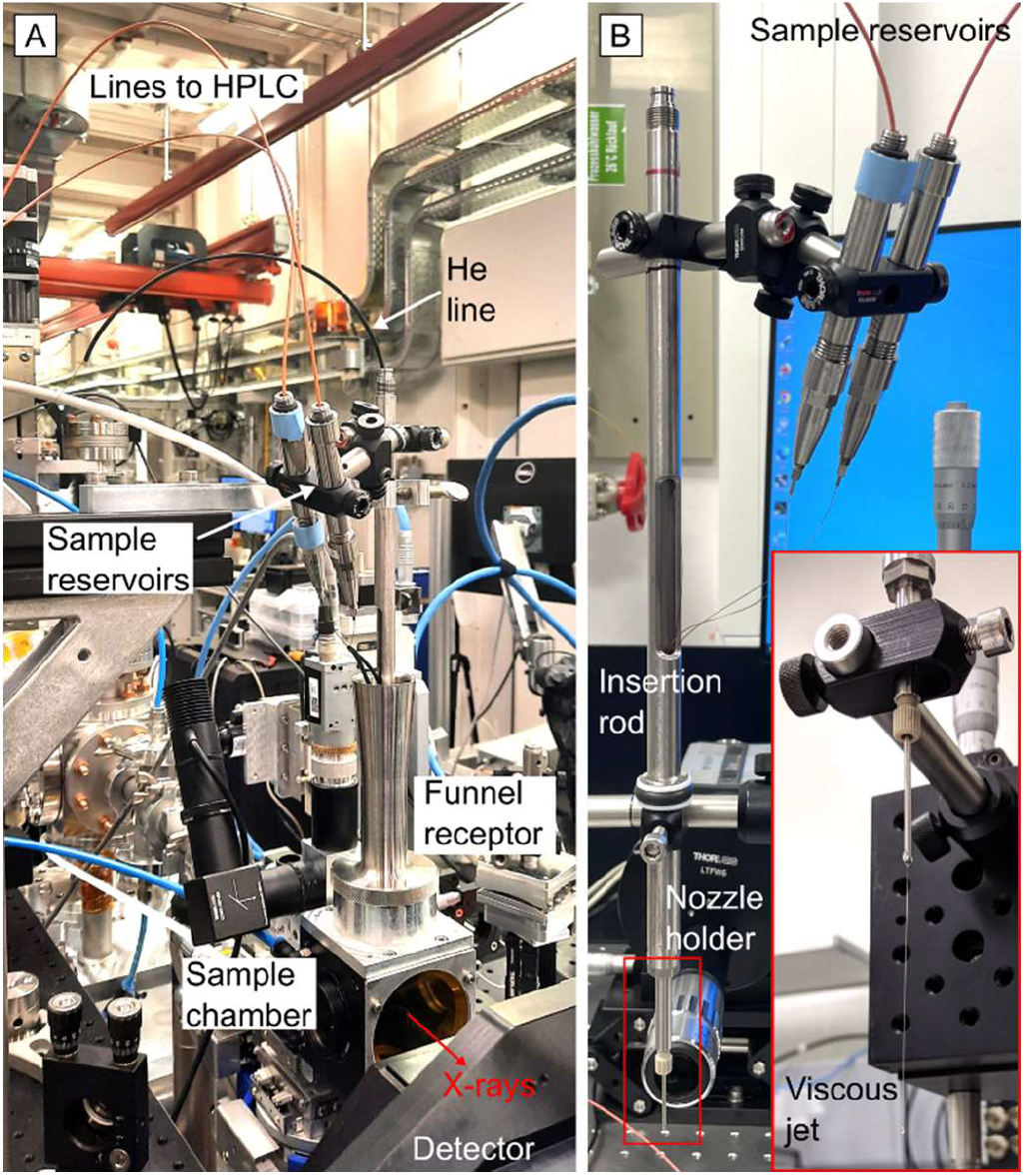
(A) Beamline injection setup surrounding the mixing-HVE device inside the helium-purged sample chamber at the SPB/SFX instrument and (B) in the laboratory test station during in-air operation.

### 2.3. Sample preparation

The preparation of iq-mEmerald protein and crystal is described in supplementary information, and the following procedures were included to prepare the samples suitable for HVE injection. The LCP was prepared with a 7:3 ratio of monoolein to water using two gas tight glass syringes and a coupler. To embed iq-mEmerald crystals (*ca*. 5×15 μm) and CuCl_2_ in LCP, 10% (v/v) crystal pellet and 20 mM CuCl_2_ solution were mixed with prepared LCP separately. For mixing investigation inside the nozzle, iq-mEmerald protein embedded in LCP was used instead of protein crystals to observe continuous fluorescence signal; 65:35 ratio of monoolein and 10 mg mL^-1^ of iq-mEmerald protein in 50 mM Tris (pH 8.0) or 10 mM CuCl_2_ was mixed and resulted in 130 μM protein and 3.5 mM CuCl_2_ in LCP, respectively (Fig. 3). The prepared samples were loaded into the HVE sample reservoirs and prepared for injection (Han *et al*., 2021).

**Figure 3.**
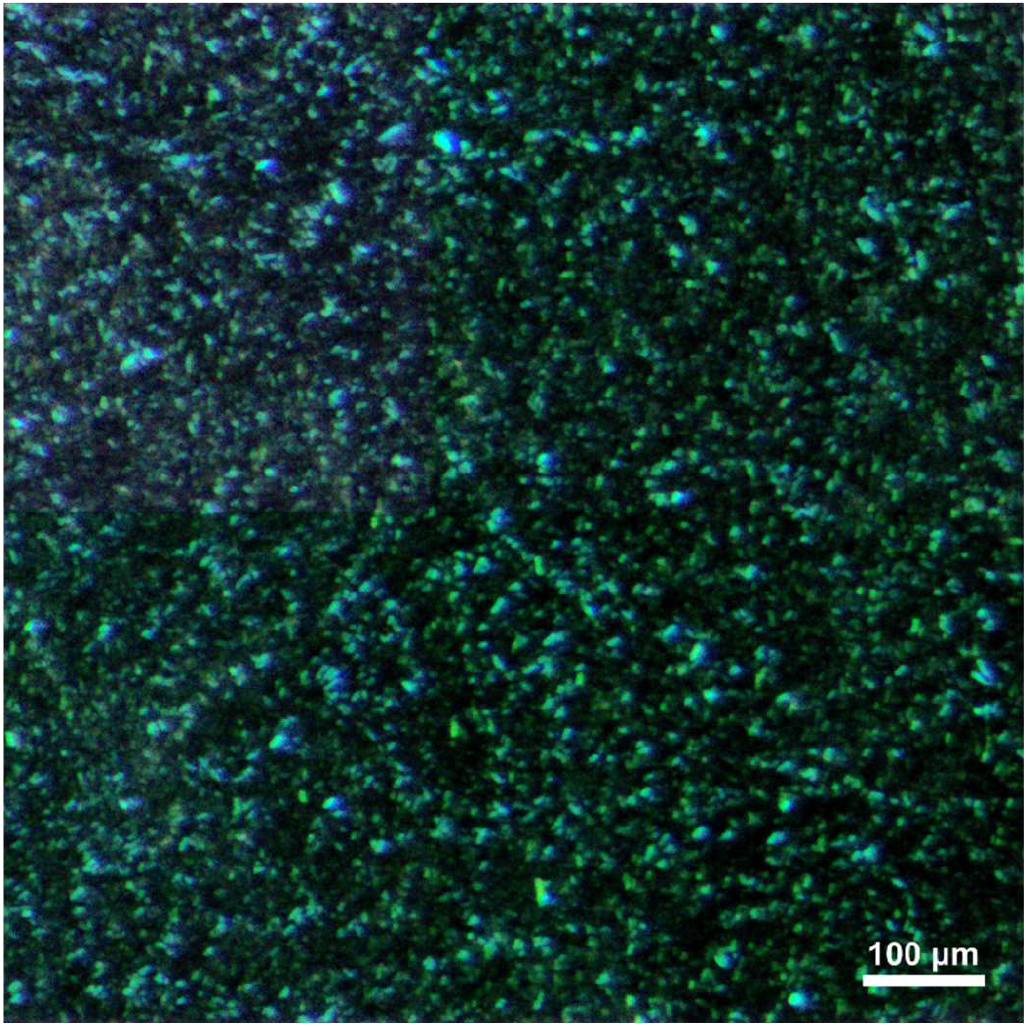
Stereomicroscopy image of iq-mEmerald crystals (*ca*. 5×15 μm) embedded in LCP, used for the mix-and-extrude device characterization.

The samples were extruded using two HPLC pumps (Shimadzu, LC-20AD XR) connected to the two HVE injector systems (Weierstall *et al*., 2014). The pumps were first run rapidly to fill the capillary with samples (up to 2.14 μL min^-1^). When the extruded sample volume was determined to be around 13 μL, shortly before the samples reached the nozzle, the flow rates were reduced to the desired values.

## 3. Results and Discussion

### 3.1. Device characterization

We investigated the fluorescence quenching of iq-mEmerald protein in LCP mixed with copper ions for the mix-and-extrude nozzle characterization on account of its facile visualization by means of optical microscopy. Moreover, the reaction time is so fast that it can be ignored in the mixing time scale of our nozzle, and the diffusion time of copper ions is expected to be fast compared with other ligands. The mixing ratio of crystal and copper ions was fixed to 1:3 so that the concentration of copper does not affect the quenching time and fluorescence intensity. The retention time for the characterization was triggered between 2.3 and 7.6 seconds (Fig. S4).

The same injection test was performed under a fluorescence microscope for quenching observation in iq-mEmerald crystals in LCP at high resolution (Movie S1). Independent of the retention time, we could observe fluorescence quenching of mixed species within the jet, meaning that the diffusion of copper ions in LCP and the quenching reaction in iq-mEmerald crystals happened faster than 2.3 seconds. The diffusion and the reaction of the test system was very fast thus the fluorescence signal disappeared even before the sample was extruded from the nozzle. To measure how fast this diffusion and reaction happens within the mixing device, we reduced the intrinsic fluorescence of the IP-S photoresist by curing the device in a UV (*λ* = 385 nm) chamber (XYZprinting, 3UD10XEU01K) for 1 hour followed by hard-baking at 80 °C for 3 hours.

Figure 4D depicts the spatial/temporal evolution of the fluorescence signal at four downstream positions: Inlet, Mixer, Nozzle and Jet. Each plot shows the extracted grey value intensity from the 2D fluorescence microscopy image as function of the channel width (indicated with yellow dashed lines in Figure 4A-C). At the ‘Inlet’, where the two liquid channels overlap, the mixing time equals 0, hence, the mixture (blue) still has the same signal as pure protein (red). The fluorescence signal at the ‘Mixer’ position, that corresponds to a mixing time of 1.8 s (based on the travelled distance of 1.02 mm), has decreased to ca. 10% of the initial 130 μM concentration at the t_0_ position and further downstream, as more time passes, there is no significant fluorescence signal inside the nozzle tip or in the free-flowing jet indicating full quenching upon mixing.

**Figure 4.**
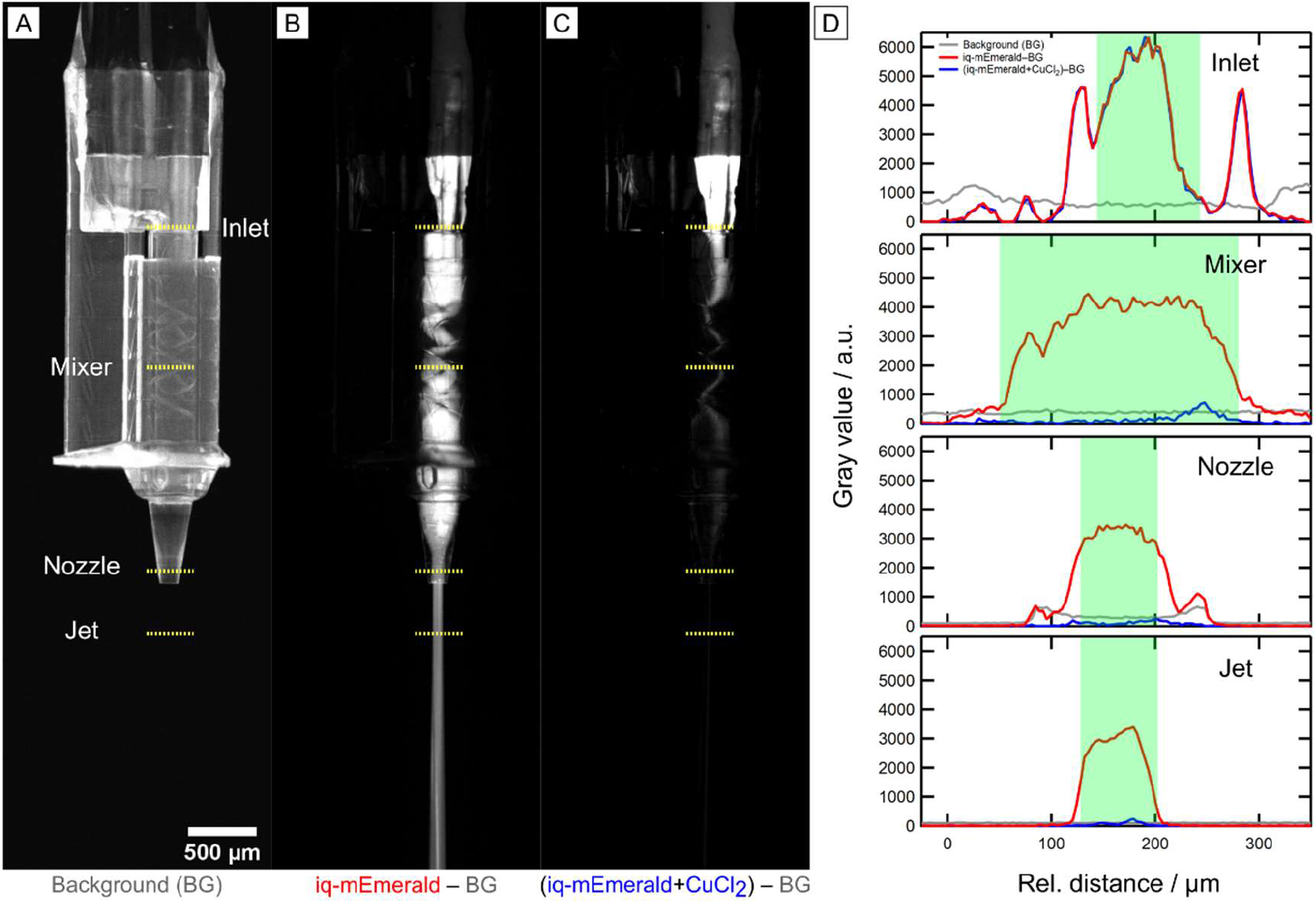
Observation of fluorescence quenching inside the 3D printed mixing-HVE device. (A) Fluorescence microscopy image of the empty device which serves as the background image. (B) Background-corrected image of the sample, *i*.*e*. iq-mEmerald protein (130 μM) in LCP flowing at 0.36 μL min^-1^ inside the device. (C) Background-corrected image showing the mixing of iq-mEmerald (130 μM) and CuCl_2_ (3.5 mM) entering with a 1:3 flow rate ratio (0.36 and 1.07 μL min^-1^, respectively). The resulting flow velocity of the extruded mixture is 5.4 mm s^-1^ and the total retention time amounts to *ca*. 4.4 s. (D) Pixel intensity profiles from the 16-bit fluorescence images (relative grey values extracted via ImageJ 1.53k) for four downstream positions: inside the 100 μm wide sample orifice (Inlet), inside the 231.7 μm wide mixing channel within the Kenics section, 1 mm downstream from the inlet (Mixer), within the final, 75 μm wide section of the device material before extrusion (Nozzle) and in-air, 300 μm away from the nozzle tip (Jet). The line positions are denoted with the yellow-dotted lines in (A). The green boxes in (D) indicate the geometric restrictions of the device/jet, which are the channel widths and jet diameter, respectively.

### 3.2. Diffusion in LCP

One of the biggest challenges in time-resolved serial crystallography, which determines distinct intermediate structures throughout the reaction, remains the rapid mixing of microcrystal with ligands (Brändén & Neutze, 2021; Schmidt, 2013). Even in investigations using aqueous solutions, the diffusion time of ligand to the crystal’s center varies based on crystal size and packing, *i*.*e*. crystals in smaller sizes or having a higher water content per unit cell are unquestionably better candidates for time-resolved crystallography, as the ligand diffusion time would be decreased for such crystals. For membrane protein crystals in LCP, diffusion is even more challenging. LCP has a similar packing structure to crystals, with the exception of a wider water channel than protein crystals. In addition, the inner diameter of our device’s mixing channel is 231.7 μm, which increases the diffusion time by 3.1 seconds for oxygen and 13.2 seconds for glucose by calculation if the ligands are assumed to diffuse directly into the center of the LCP stream (Atkins & Paula, 2006). To reduce such diffusion time caused by the width of the stream, a Kenics architecture was employed in our mixing channel to support the mixing of crystal and ligand in LCP. With the series of helical elements for repeated flow splitting/stretching, we exponentially increase the diffusive interface with each additional element. Mixing time point uniformity/dispersion becomes more dependent on spatial location along the mixer and highly insensitive to flow rate fluctuations. This feature is in contrast to flow-focusing designs that tune diffusion distances through flow rate differentials. Even with these efforts to obtain discrete intermediates by reducing diffusion time, it is still likely to observe a mixture of multiple intermediates by mixing with this device. With current advancements in data analysis (*e*.*g*. Xtrapol8, (De Zitter *et al*., 2022)), it is projected that mixed diffraction data of multiple states can be separated.

Due to the complexity of the LCP structure and membrane protein, it is difficult to predict the diffusion and reaction time of ligand binding in LCP; for instance, if the ligand is diffused in an aqueous channel and the binding site of membrane protein is located in an aqueous channel, the diffusion time would be comparable to that of in aqueous solution. Alternatively, if the binding occurs in the lipid, it is likely too time-consuming to be studied using this type of device. Therefore, we assume that this mixing-HVE is suited for the examination of reactions taking place on the cell/membrane surface, where most current drugs targeting membrane protein act on (Yin & Flynn, 2016). However, due to this overall unpredictability of reaction in LCP, it is highly recommended to characterize the reactions in LCP prior to X-ray diffraction using different biochemical and biophysical techniques in order to confirm that the reaction takes place on a timescale compatible to the operation of these nozzles.

### 3.3. Time-resolved X-ray diffraction

To demonstrate the device’s performance under real-life serial membrane protein crystallography conditions/requirement, we collected pre/post-mixing diffraction patterns of GPCR (G protein-coupled receptor) membrane protein crystals in LCP at the SPB/SFX instrument of the European XFEL using a photon energy of 12.4 keV, an X-ray pulse repetition rate of 10 Hz and a beam size of ca. 2×2 μm. The pulse energy was 2.2 mJ and the pulse duration was *ca*. 100 fs. The detector (JUNGFRAU 4M) was positioned 120 mm away from the sample stream.

For injection, a co-flowing sheath of helium gas was used and the supply pressure was adjusted to 140 psi (corresponding to 24 mg min^-1^) to maintain a stable sample flow. The distance from nozzle tip to X-ray interaction was *ca*. 200 μm. For the *in situ* mixing, a combined flow rate of 0.36 μL min^-1^, *i*.*e*. 0.11 μL min^-1^ (sample in LCP) + 0.25 μL min^-1^ (ligand in LCP), with associated pressures of *ca*. 450 psi and 420 psi at the HPLCs, was applied. This led to a flow velocity of 1.35 mm s^-1^ for the extruded mixture (sample width = 75 μm) and a probed time point of *ca*. 18.5 s. The un-mixed sample was pumped with a flow rate of 0.36 μL min^-1^ (*i*.*e*. 1.35 mm s^-1^) with an associated pressure of *ca*. 480 psi at the HPLC. Figure 5 shows exemplary diffraction patterns showing strong reflections from the crystals in both states. Details on the sample and data statistics are currently being prepared for a forthcoming article.

**Figure 5.**
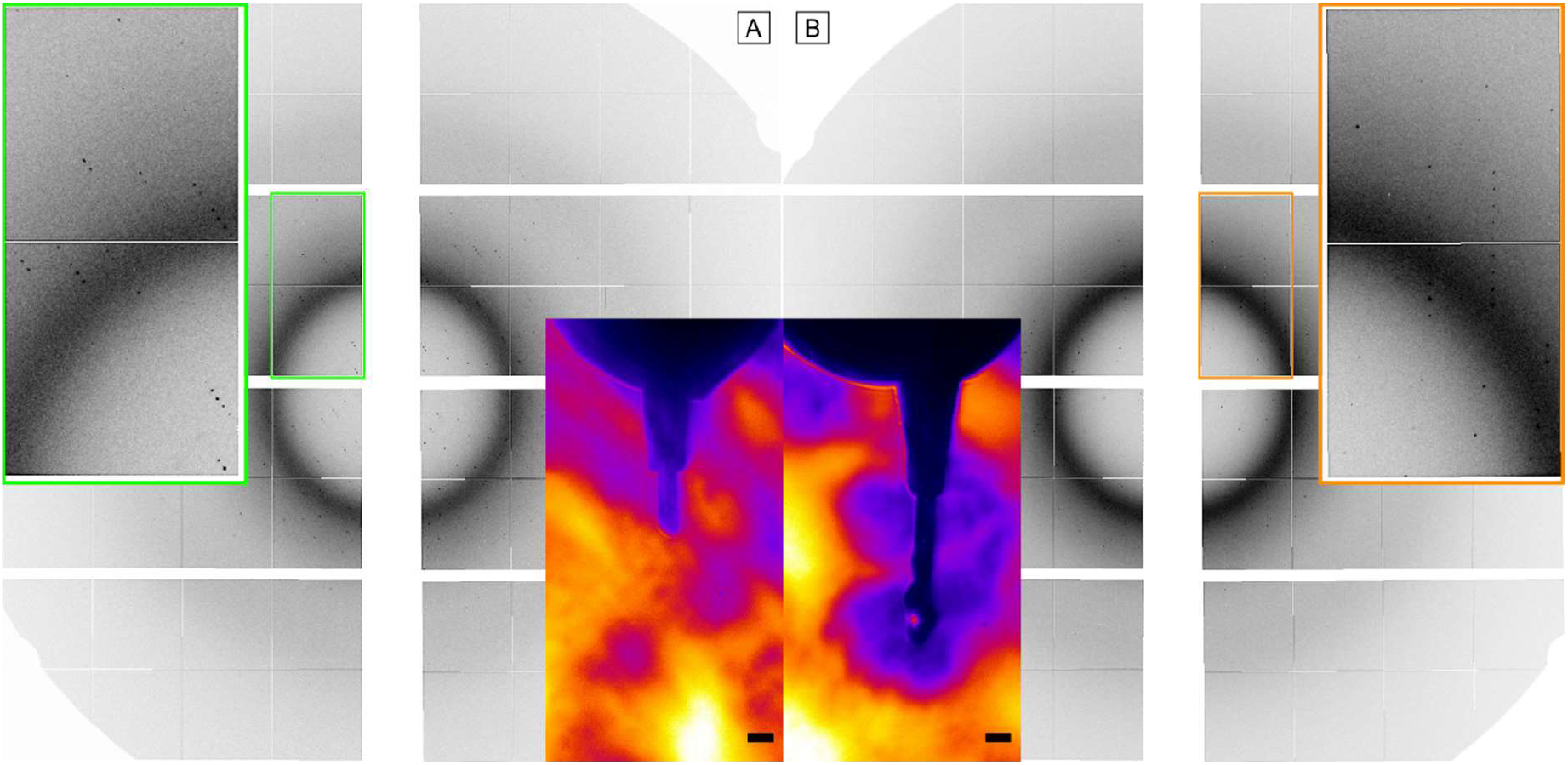
Diffraction patterns of membrane protein crystals (A) without mixing and (B) after mixing with ligand (*t* ∼18.5 s) collected with the JUNGFRAU 4M detector (pixel size is 75×75 μm^2^) at room temperature. The detector consists of 8 front-end modules, each being 80×40 mm (1024×512 pixels) large. The prominent ring arises from monoolein scattering at a 4.5 Å resolution. In the inlay, the beamline side-view microscopy images are showing the respective viscous extrusion from the nozzle tip (the scale bar denotes 100 μm.).

### 3.4. Outlook

The current setup, which utilizes externally positioned fluid-feeding capillaries, can be susceptible to capillary rupture. Moreover, sample-conserving inner diameters below the here used 250 μm can be prone to generate high pressures during injection. It is also worth noting that the used capillary length of 30 cm generated a void volume of 14.7 μL. Especially when considering the difficulties of membrane protein sample preparation and the available sample reservoir capacities of 40 and 120 μL, this sample loss might not be negligible. To reduce the length of the capillaries, a more compact mixing-HVE composed of two hydraulic systems within a single, protective injector rod is desired. In addition, the installation of injection infrastructure with a large footprint should be avoided to not interfere with other significant instrumentation (detector, laser setup, electronics, optics, etc.) already present at the beamline.

The design of another type of HVE described in the recent publication by Shimazu *et al*. (Shimazu *et al*., 2019), allows a capillary connection to the sample reservoir without the need of direct screwing. Thus, we assume that an adaptation of their design would enable the advancement of a compact mixing-HVE utilizing very short capillaries.

As described above, diffusion time of diluted species in LCP media is difficult to predict. To cover a broader range of diffusion and reaction time, modification on the device design is under consideration. For shorter retention time, the length of the mixer can be reduced. To access longer retention times, a modular assembly approach can be pursued in which the 3D printed mixer and nozzle are connected by a capillary extension of custom length. In addition, once the compact mixing-HVE is commissioned and shorter capillaries can be used, the ID of capillaries and mixer/nozzle can be reduced because pressure build-up due to long capillaries is not a concern anymore. Then, even higher sample flow rates can be applied to the system: with faster sample movement in the mixer, the retention time can be reduced further.

## 4. Conclusion

We fabricated and characterized a mix-and-extrude nozzle for time-resolved protein serial crystallography. Fluorescence quenching by mixing iq-mEmerald crystals with copper ions was observed on a short timescale, indicating that the diffusion in LCP occurred in a negligible time compared to the retention time of the mixer. With current designs of the device, a retention time between *ca*. 2 and 20 seconds can be triggered, and injectors aiming for shorter and longer retention time can be prepared with minor design modification. The first EuXFEL user experiment using this nozzle in a helium environment at atmospheric pressure has been conducted and the manuscript describing the result is currently in preparation. We anticipate that this mix-and-extrude nozzle will be widely used for mixing experiments of viscous media containing, for instance, membrane protein crystals at synchrotrons and FELs, hence, efforts to create a mixing-HVE as a single compact system are ongoing.

### 5. Related literature

The following references, not cited in the main body of the paper, are cited in the supporting information: Samarkina *et al*. (2009); Yu *et al*. (2014).

## Acknowledgements

The authors would like to thank all members of the EuXFEL SEC group for the excellent teamwork, their technical support and helpful discussions. We further acknowledge the EuXFEL in Schenefeld, Germany, for provision of XFEL beam time at SPB/SFX and would like to thank the staff for their assistance. We also thank Thomas Dietze and Marco Schrage (EuXFEL) for CNC machining. Moreover, we express our gratitude to Richard Neutze (University of Gothenburg), Juraj Knoška (Center for Free-Electron Laser Science, Hamburg) and Michael Heymann (University of Stuttgart) for fruitful discussions.

Tiankun Zhou was supported by a Wellcome Investigator Award (210734/Z/18/Z; to Allen M. Orville, Diamond Light Source).

## Supporting information

### S1. Materials and Methods

#### S1.1. Device fabrication and assembly

The 2PP-based microfabrication followed general guidelines detailed elsewhere (Knoška *et al*., 2020). 3D geometries were designed in AutoCAD (Autodesk) or NX (Siemens) and exported as STL formats.

Using DeScribe (Nanoscribe), the STL-based 3D designs were converted to print-job instructions (GWL). For fast printing times, slicing distances of 2 μm, hatching distances of 0.7 μm, and block sizes of 285/285/299 μm (x/y/z) with 15° block shear angles, 2 μm block overlaps and 1 μm layer overlaps were chosen.

The devices (solid volumes) were then printed with the IP-S photoresist using the Nanoscribe Photonic Professional GT equipped with a 25× objective lens (Zeiss) in upwards direction (+z) with alternating hatch lines and in the dip-in mode. IP-S was deposited onto an indium tin oxide (ITO) coated glass slide. The laser power was 100% (*i*.*e*. 156 mW exiting the laser source, 70 mW arriving at the objective), and print speeds were 100,000 μm s^-1^. With these parameters, the printing time for one mixing-HVE device was 3 h for J_7 and 2 h for J_8, respectively. It is worth noting that the shorter design allows the printing of multiple devices in one batch (*i*.*e*. one glass slide) which reduces the overall device fabrication time.

For the device assembly, two fused silica capillaries (Polymicro, OD 360 μm, ID 250 μm), each 30 cm in length, were inserted into the liquid access ports of the 3D printed IP-S device and glued with freshly mixed (low-viscous) epoxy glue (Devcon 5-minute epoxy for general purpose) (Fig. 1). It was crucial that the amount of applied glue does not exceed the total width of the 2PP–3D printed part. Otherwise, the steel tubing cannot be imposed on the IP-S–capillary assembly. To minimize sample waste due to lengthy capillaries, one might consider the connecting of a cartridge with pure LCP after the sample is run empty to push out the remaining crystal from the capillary.

After epoxy curing for 12 h at RT, the two capillaries were run through a steel tubing with 0.046 in ID (IDEX, part no. U-145), which was brought into close proximity to the ‘collar’ of the IP-S tip at the other end. A small gap of the width of a capillary was left between steel tubing and the nozzle’s ‘collar’ to allow the deposition of a small amount of slightly viscous epoxy glue (cured for 2 min). After thus bridging the steel and IP-S with strong epoxy bonds, the steel tubing was connected to a T-union (P-728 or custom variant) using a #10-32 UNF (IDEX, part no. F-333N) fitting. The through-hole of the T-union must be larger than 1 mm to allow room for two capillaries. The opposite port of the T-union (port #2, *i*.*e*. where the two capillaries exit) allows connection at the Interaction Region Downstream (IRD) at the SPB/SFX instrument via insertion rod (Fig. S1A&B). (Round *et al*., in preparation) Alternatively, mounting can occur using the nozzle holder (Weierstall, 2014) and standard mounting parts (Fig. S1C). The exiting fluid-feeding capillaries can then be connected to either pressure-driven syringe pumps or the ASU injector system for pumping high-viscous samples (Weierstall *et al*., 2014).

The third port of the T-union is used to connect another capillary for introducing pressurized helium using a flexible tubing sleeve (F-242, 0.0155 in ID) in combination with a F-354 nut and a LT-135 ferrule (all from IDEX). In this manner, the helium enters the steel tubing and runs through the 3D printed pockets of the injector to sheath the mixed liquid streams after extrusion (Fig. 2A). To make sure that the helium is not released through the second port, a dual-lumen sleeve (FEP MultiLumen, Zeus Industrial Products, Inc.) can be used as shown in Fig S1A. Alternatively, the third capillary for helium supply can be simply inserted directly into the back of the steel tubing (where the two liquid-feeding capillaries exit) and sealed with epoxy glue. Details on utilized injection hardware can be found elsewhere (Vakili *et al*., 2022).

**Figure S1.**
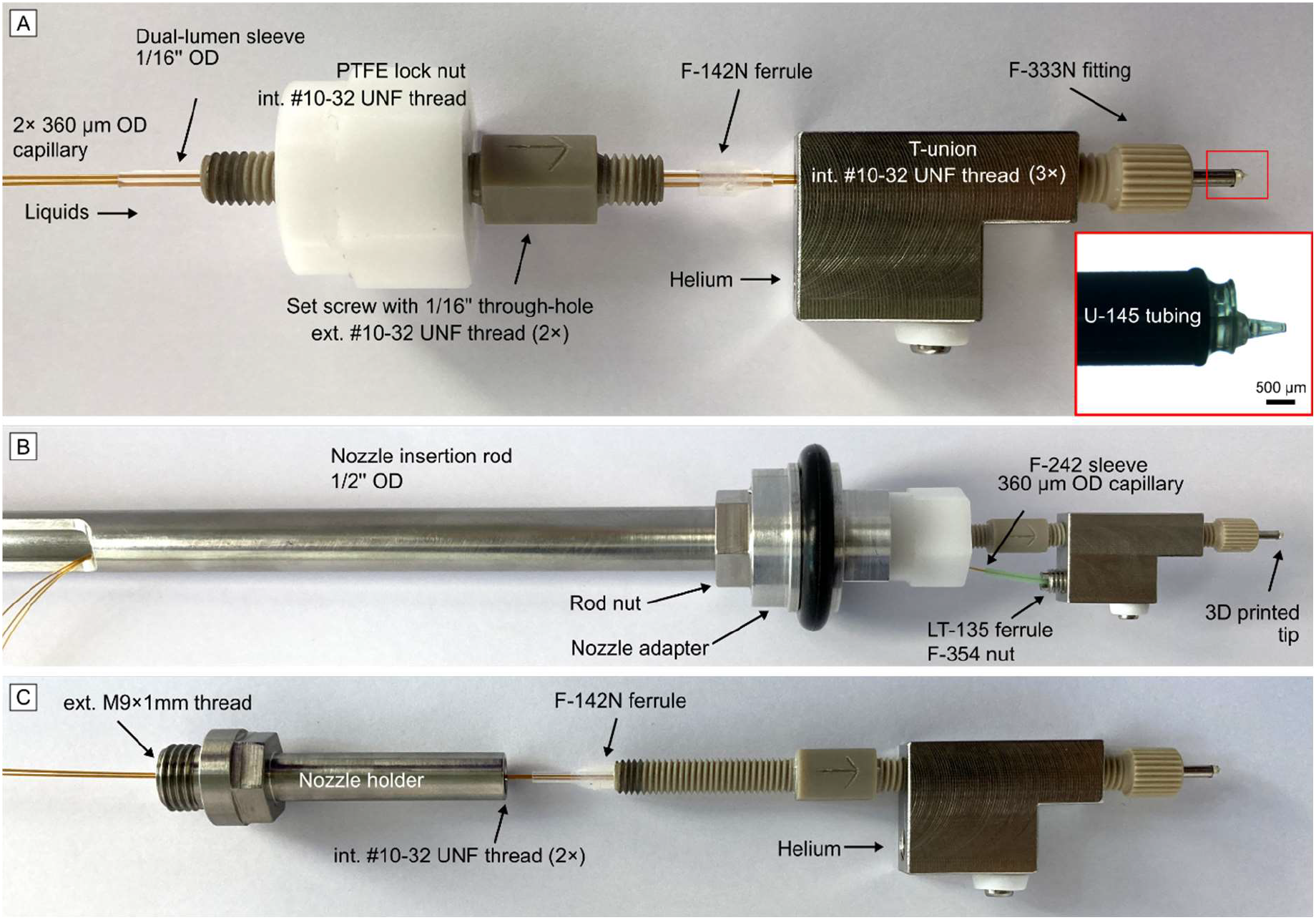
Photographs of the injector assembly surrounding the 3D printed mixing-HVE device. The use of wider ID capillaries (250 μm), allows the injector tip to be in ca. 30 cm distance from the feeding sample reservoirs (not depicted here). (A) Close-up of the injector system connected to a custom steel T-union with description of utilised fluid connection parts. For a uniform helium stream, which enters the third port of the T-union and exits through the injector tip, a dual-lumen sleeve, which encases two liquid capillaries at once, is used. (B) Depiction of the injector system after connection to a 30 cm long insertion rod for in-vacuum/in-helium beamline installations at the IRD@SPB/SFX. The connective nozzle adapter contains an internal #10-32 UNF thread for device mounting and an external groove for a 25×5.5mm O-ring for sealing the sample chamber. (C) Instead of using the nozzle adapter, a combination of a dual-lumen sleeve, dual-fittings (#10-32 UNF threads) and the ‘nozzle holder’ can be used for in-air operation. Here, at the right end of the nozzle holder, the tan dual-fitting is inserted and the assembly continues as shown in (B).

#### S1.2. Preparation of iq-mEmerald microcrystals

Iq-mEmerald is a GFP-derivative designed and studied by Yu *et al*. (Yu *et al*., 2014). It has an UV-Vis absorption maximum at 488 nm and an emission maximum at 512 nm. The pET17b-iq-mEmerald expression vector was purchased from Biocat GmbH (Heidelberg, Germany). One colony of *Escherichia coli* BL21 DE3 pET17b-iq-mEmerald was transferred into 50 mL LB containing 100 μg mL^-1^ Ampicillin and was incubated at 37 °C for 16 h at 180 rpm. The next day, the OD_600nm_ was adjusted to 0.1 for 1 L LB containing 100 μg mL^-1^ Ampicillin and incubated at 37 °C and 180 rpm until the OD_600nm_ reached 0.6.

Recombinant expression of iq-mEmerald was induced by adding 0.5 mM IPTG. Cultures were cultivated for 16 h at 18 °C and 180 rpm. Cells were harvested at 8000× g for 1 h at 4 °C and cell pellets were stored at -80 °C. The purification was adapted from Samarkina *et al*. (Samarkina *et al*., 2009). Cell lysis was performed after resuspending the pellet in lysis buffer (20 mM Tris, pH 7.8, 150 mM NaCl) using a sonicator. After centrifugation for 15 min at RT and 11000 rpm, the supernatant was heated up for 15 min at 65 °C. Reaction tubes were spun down for 15 min at 11,000 rpm and the supernatant was collected. 10 mL of cell lysate was rapidly mixed with 3 mL of 5M NaCl and 23.3 mL saturated (NH_4_)_2_SO_4_ (pH 7.8). 12 mL of 96% Ethanol was added instantly and vortexed for 30 sec. Samples were spun down for 7 min at RT with 3000× g. Iq-mEmerald is present in the organic phase which is carefully removed and diluted to 20% (NH_4_)_2_SO_4_ saturated solution in 20 mM Tris, pH 7.8. The sample was filtered with a 0.2 μm filter and subjected to a HIC (hydrophobic interaction chromatography) column that was equilibrated with 20 mM Tris (pH 7.8), 20% (NH_4_)_2_SO_4_ saturation. Sample was eluted over 20 column volumes of elution buffer (20 mM Tris, pH 7.8). The purified protein was concentrated to 50 mg mL^-1^.

For batch crystallization, 500 μL protein solution (50 mg mL^-1^) were mixed with 500 μL of 3 M (NH_4_)_2_SO_4_, 50 mM Tris (pH 8.0) as well as 5 μL seed-stock and vortexed for 30 sec. After 1 min, additional 500 μL of 2.5 M (NH_4_)_2_SO_4_, 50 mM Tris (pH 8.0) were added and vortexed briefly. Crystals grew overnight to 15 μm×5 μm. Crystals were filtered with a 20 μm gravity filter before being embedded in LCP.

#### S1.3. Injection test station with fluorescence microscopy setup

Fluorescence microscopy was conducted with a Zyla 4.2 sCMOS camera equipped with a 2× objective (Mitutoyo, #46-142) leading to a pixel size of 3.6 μm. As a light source, a fiber-coupled LED (ThorLabs, M490F3), *λ* = 490 nm (FWHM = 26 nm), together with a fluorescence imaging filter set, composed of a 475/35 nm FITC excitation filter (MF475-35) and a 525/39 nm GFP emission filter (MF525-39), has been used (Fig. S3).

**Figure S2.**
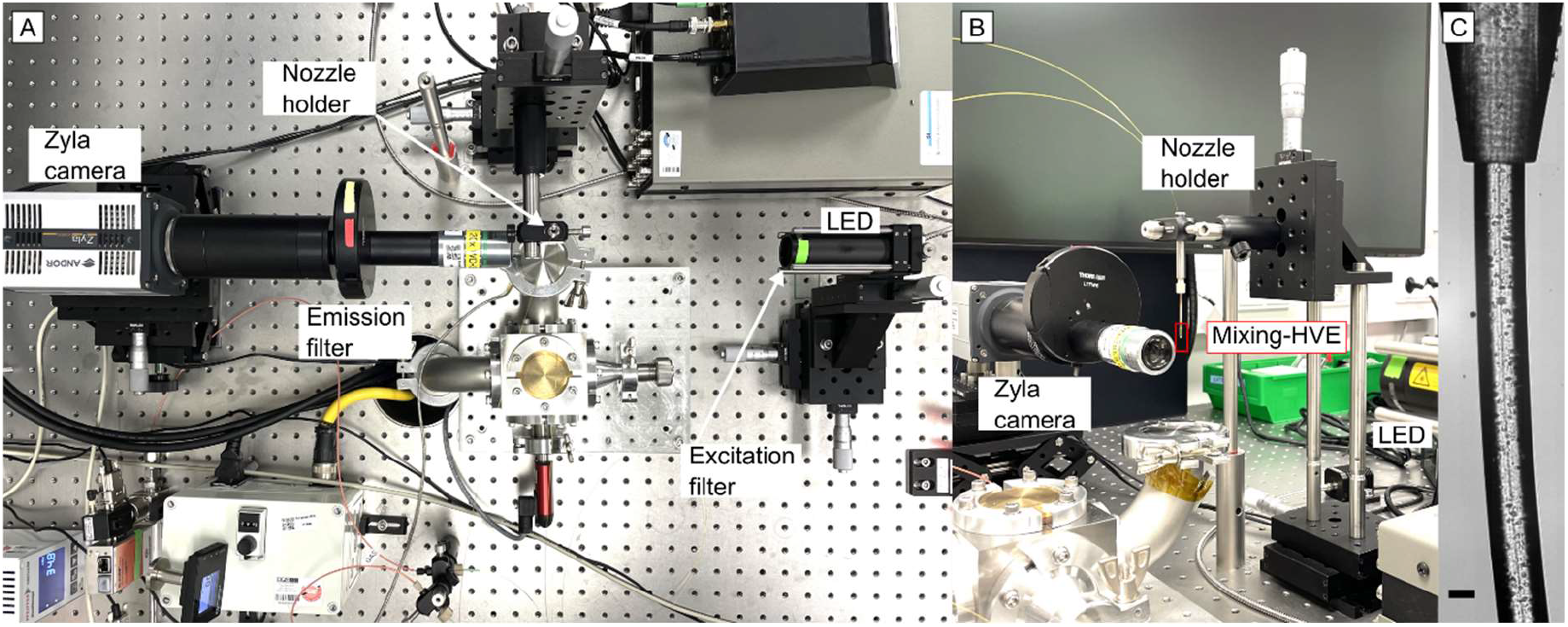
Injection test station setup from (A) the top view and (B) from the side view depicting the LED light source with excitation filter and Zyla camera with emission filter. (C) Detailed view of the mixing-HVE tip (here with a 100 μm ID). The exemplary microscopy image (125×615 pixels, pixel size ca. 1.9 μm) depicts the dual-extrusion of LCP media, each delivering *ca*. 5 μm wide lysozyme crystals with a total flow velocity of 1.3 mm s^-1^ (as used for detector calibration). The scale bar denotes 50 μm.

**Figure S3.**
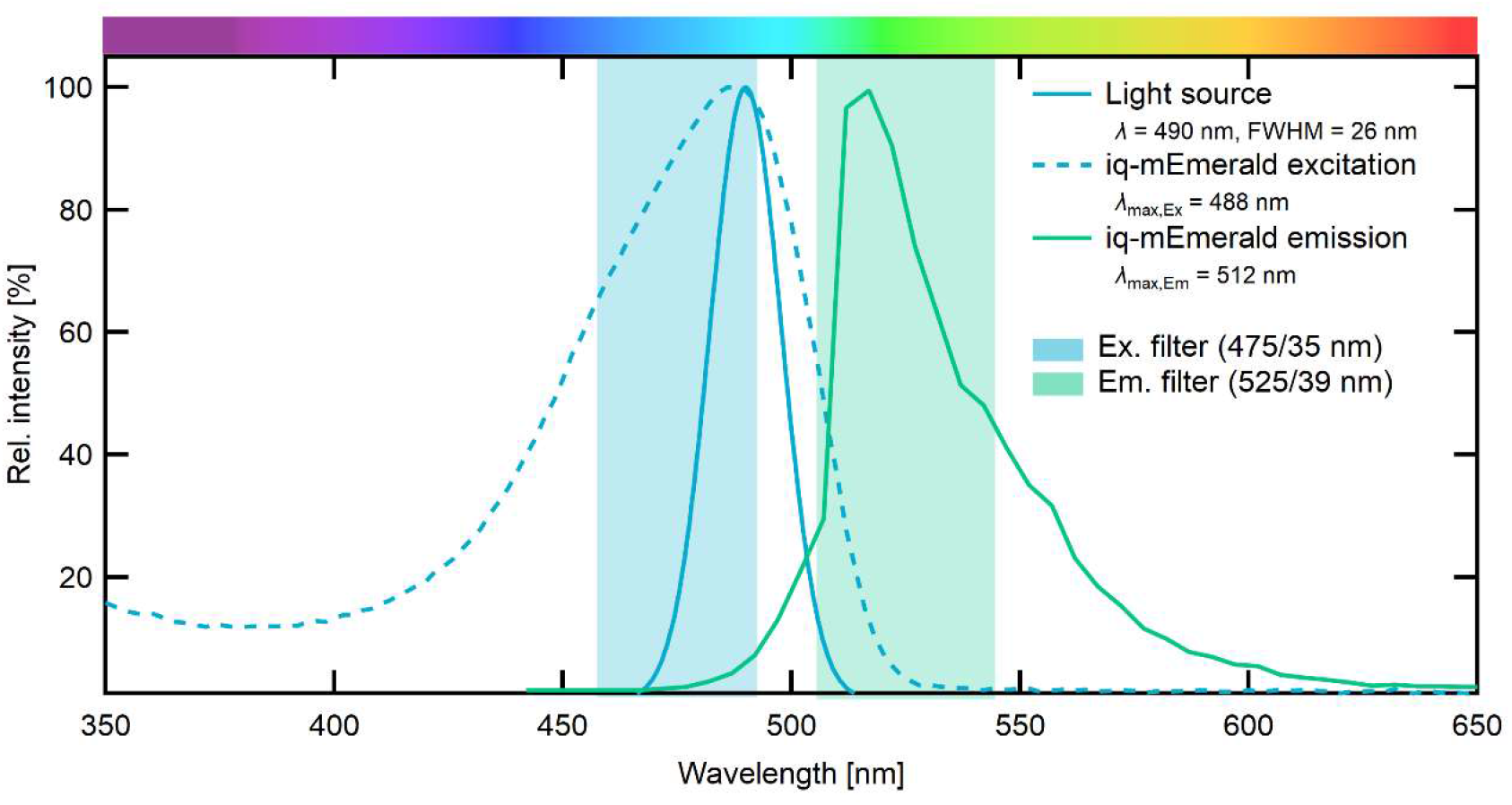
UV-Vis absorbance and emission spectra for the iq-mEmerald crystals with indicated excitation/emission filter range of the imaging system (see S1.3.) and spectral line of the 490 nm (FWHM = 26 nm) LED light source. The absorption spectrum of the protein in solution was recorded with a Tecan Spark 10M microplate reader. The emission spectrum was acquired from a single crystal in 5 nm steps on a Nikon AX-R confocal microscope equipping a DUX-VB4 detector, using the 405, 445, and 488 nm channels (LUA-S6 laser unit) for excitation and the spectral acquisition mode from the Nikon NIS-Elements AR 5.41.01 software.

**Figure S4.**
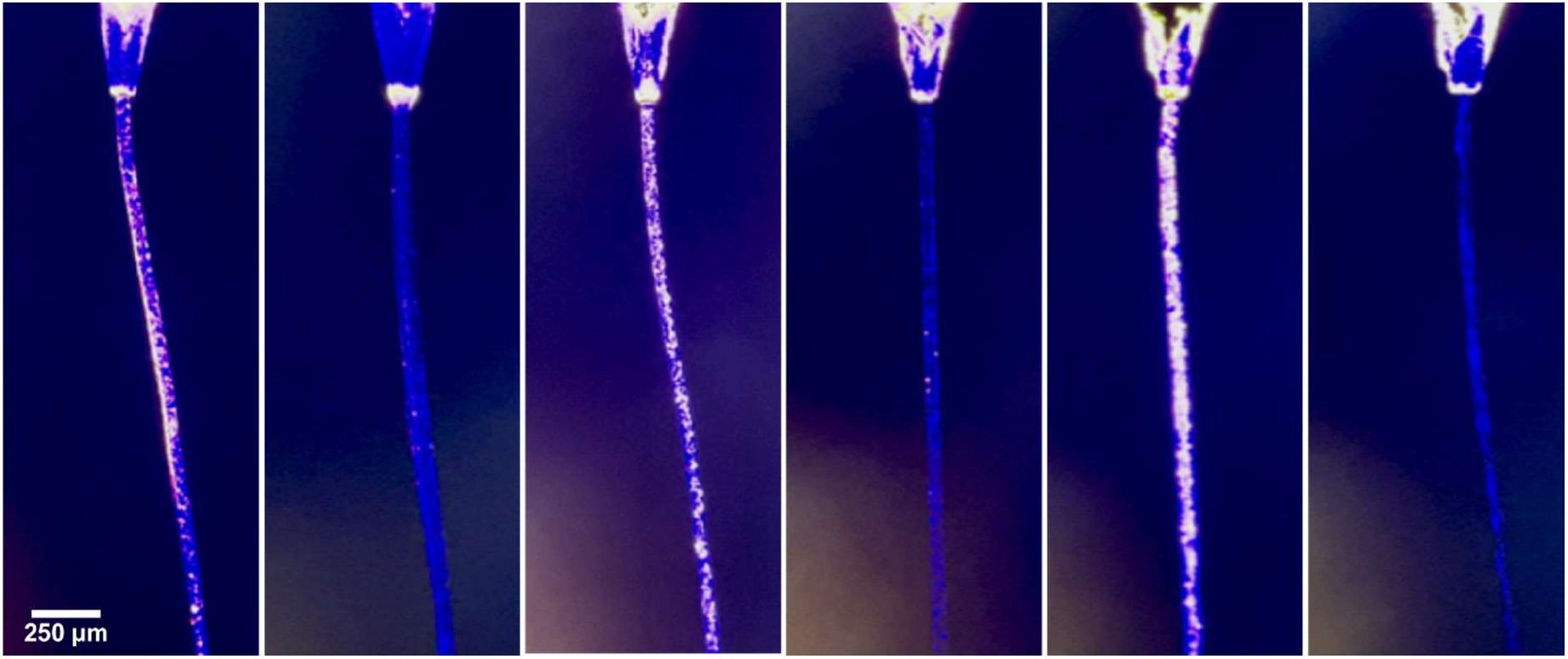
Fluorescence microscopy images taken at the setup described in S1.3. depicting (left) the extrusion of iq-mEmerald crystals (*ca*. 5×15 μm) and (right) fluorescence quenching of iq-mEmerald crystals by mixing with Cu^2+^ with variation of the flow velocity (thus retention time in the mixer) (22.5, 4.5, and 2.2 seconds, respectively).

#### S1.4. Fluorescence videography setup

12-bit frames (2048×1048 pixels, 2fps) were recorded on a Nikon AX-R confocal microscope equipped with a Plan Apo lambda 20× objective (NA = 0.75), using the resonant bidirectional scanner mode and a pinhole size set to 34.9 μm. A 488 nm laser was used for excitation with an emission gate between 499 nm and 551 nm. Image sequence was processed to reduce the shot noise from the resonant scanner using the deep learning-based Denoise.ai algorithm from the Nikon NIS-Elements AR 5.41.01 software, background-corrected and exported to an 8-Bit RGB compressed MP4 video file.

**Movie S1.**
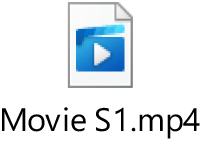
Fluorescence microscopy video (see S1.4.) showing fluorescent iq-mEmerald crystals in LCP while being extruded from the tip of the nozzle and subsequently quenched by flow start of CuCl_2_.

